# Network depth affects inference of gene sets from bacterial transcriptomes using denoising autoencoders

**DOI:** 10.1101/2023.05.30.542622

**Authors:** Willow Kion-Crosby, Lars Barquist

## Abstract

The increasing number of publicly available bacterial gene expression data sets provides an unprecedented resource for the study of gene regulation in diverse conditions, but emphasizes the need for self-supervised methods for the automated generation of new hypotheses. One approach for inferring coordinated regulation from bacterial expression data is through the use of neural networks known as denoising autoencoders (DAEs), which encode large datasets in a reduced bottleneck layer. We have generalized this application of DAEs to include deep networks and explore the effects of network architecture on gene set inference using deep learning. We developed a DAE-based pipeline to extract gene sets from a large compendium of transcriptomic data in *Escherichia coli*, independently of the DAE network parameters and architecture. We validate our method by identifying many of the inferred gene sets with known pathways in *E. coli*, and have subsequently used this pipeline to explore how the choice of network architecture impacts gene sets recovery. We find that increasing network depth leads the DAEs to explain gene expression in terms of fewer, more concisely defined gene sets, and that adjusting the network compression results in a trade-off between generalizability and overall biological inference. Finally, leveraging our understanding of the impact of DAE architecture choices on gene set inference, we apply our pipeline to an independent uropathogenic *E. coli* dataset collected directly from infected patients to identify genes which are uniquely induced during human colonization.

## Introduction

The regulation of gene expression is central to how bacteria respond to changes in their environment. Changes in the expression of just a few genes can activate systems designed for the uptake of nutrients (Davies *et al*., 2021; Schalk and Guillon, 2013), lead to the assembly of flagella for rapid motility to seek out nutrient sources (Armitage and Berry, 2020), to the formation of biofilms leading to multiple resistances (de la Fuente-Núñez *et al*., 2013), or to other transcriptomic responses associated with persistence in hostile conditions or with antibiotic resistance (Seo *et al*., 2015; Gollan *et al*., 2019). Indeed, also often specific transcriptomic programs are associated with host colonization and pathogenicity (Fang *et al*., 2016). In principle, transcriptomics can provide a complete picture of gene expression in diverse conditions, and serve as a basis for the exploration of fundamental questions regarding bacterial behavior including the mechanisms underlying bacterial infection, persistence, resistance.

Inferring the regulatory networks that govern microbial gene expression has a long history. Early work relied on large collections of microarray data, with a wide variety of methods applied (Marbach *et al*., 2012). These include approaches based on regression (Bonneau *et al*., 2006, 2007), mutual information (Faith *et al*., 2007), and correlation (Langfelder and Horvath, 2008). The development of bulk RNA sequencing has added new impetus to the field with the growing availability of probe-independent transcriptomics data spurring the development of new methods (Sastry *et al*., 2019; Tan *et al*., 2020; Yuan *et al*., 2022).

Recent years have also seen the emergence of deep learning as a new analysis tool for genomics and transcriptomics (Zou *et al*., 2019). The advantages of deep learning are due to the presence of “hidden” layers between the input and output of the network. These additional layers, the number of which we refer to as the network depth, allow the models to perform complex, hierarchical data processing which leads to state-of-the-art performance on a variety of applications (LeCun *et al*., 2015; Eldan and Shamir, 2016). A particular type of neural network known as an autoencoder (AE) allows for self-supervised training by automatically learning to compress each data point into a low-dimensional-but-meaningful representation through what is called the bottleneck (BN) layer (Hinton and Salakhutdinov, 2006). Autoencoders consist of two networks, an encoder and a decoder, which are trained simultaneously. Denoising autoencoders (DAEs) differ from traditional autoencoders by the addition of randomly added artificial noise at the input which forces the DAEs to learn the true relationship between features (Vincent *et al*., 2008).

DAEs have been applied to the inference of co-expressing sets of genes from microarray data (Tan *et al*., 2017, 2016): DAEs not only have the advantage over other methods of being self-supervised, but also have the potential to uncover nonlinear relationships between genes given that they can be deep learning models. The authors used shallow DAEs, or networks where the input layer is directly connected to the BN layer. This choice allows for direct model interpretability through examination of the weights connected to each BN node. Additionally, they further developed an ensemble method (Tan *et al*., 2017) where many networks are trained simultaneously to circumvent model instability, as networks trained from different initial conditions yield different results when trained on limited data sets.

DAEs, however, are extremely flexible. The number of layers, size of each layer, and in particular, the size of the BN layer is left to the discretion of the operator. In the context of gene expression analysis, it is unclear how these architecture choices affect downstream analysis. We have developed a pipeline for the inference of gene sets from bacterial expression data using DAEs which allows for the use of both shallow and deep networks: we examine the changes in predicted gene expression at the output of the decoder when each BN node is independently activated. We have used this pipeline to investigate how the choice of DAE architecture impacts the recovery of gene sets by evaluating the performance of a variety of architectures on the PRECISE 2.0 RNA-seq compendium (Lamoureux *et al*., 2021) for *Escherichia coli*. Finally, leveraging our understanding of DAE architecture choice, we use our inference pipeline to recover new gene sets uniquely associated with human infections from a uropathogenic *E. coli* (UPEC) transcriptomic data set (Subashchandrabose *et al*., 2014). This work extends the usage of DAEs to include deep models, and illustrates the advantages of network depth in inferring coherent gene sets from transcriptomic data.

## Results

### A pipeline for autoencoder architecture exploration

To address the question of how the choice of network architecture impacts biological inference from bacterial expression data, we have trained 100 randomly initialized denoising autoencoders (DAEs) with a variety of shallow and deep network architectures. For training we used the PRECISE 2.0 *E. coli* K-12 RNA-seq compendium (Lamoureux *et al*., 2021), consisting of 815 experiments using a standardized RNA-seq protocol executed in a single lab (**Fig. 1A**). For each trained network in each ensemble, we then subsequently activate each node at the bottleneck (BN) layer and propagate this signal through the decoder. This provides a predicted expression value for each gene which we then examine to determine which biological processes these BN nodes represent.

**Figure 1.**
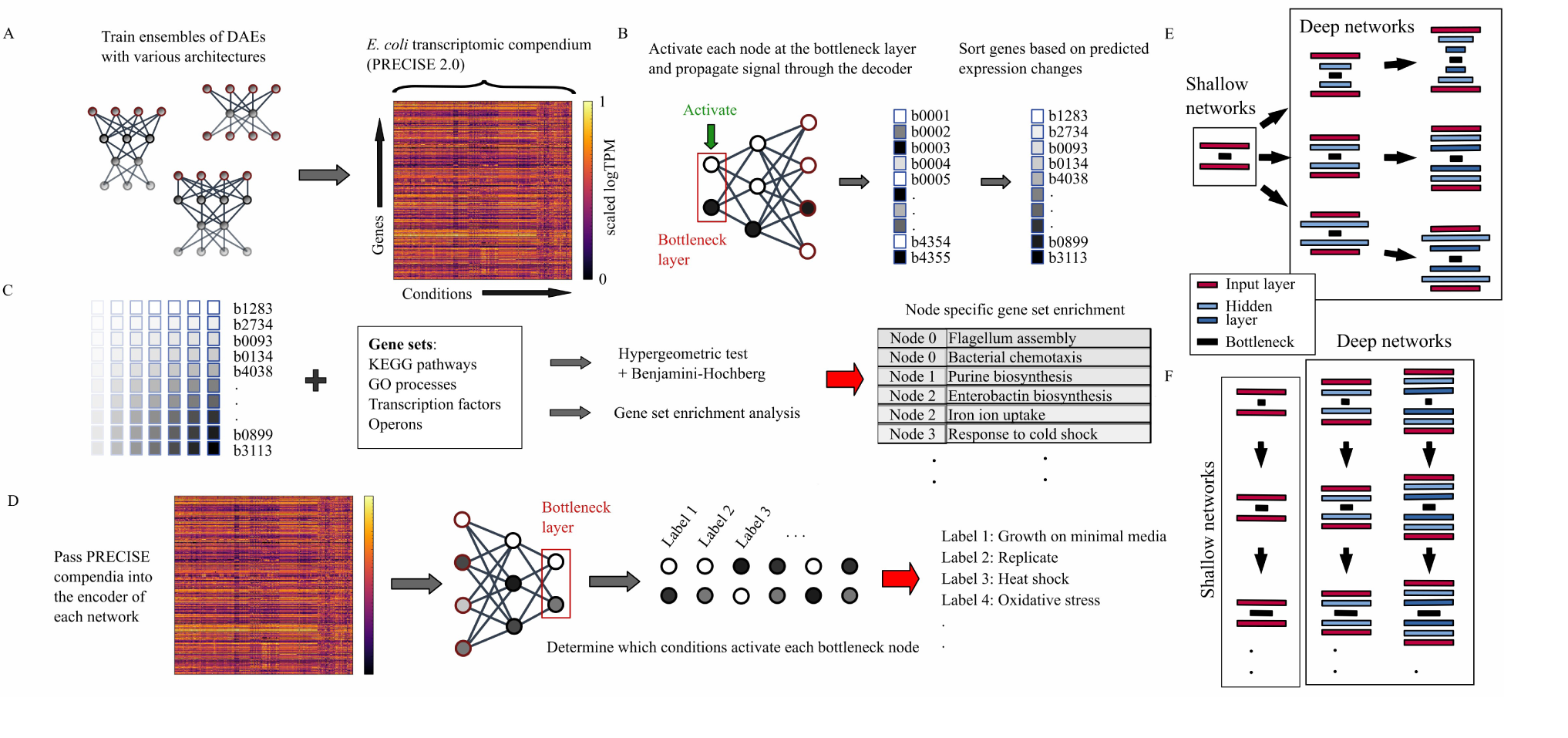
Schematic of DAE architecture exploration pipeline. (A) Each DAE architecture ensemble is trained on the PRECISE 2.0 gene expression compendium. (B) The bottleneck nodes of each architecture are subsequently activated, and the activations are propagated through the decoder. These gene BN vectors are then sorted such that genes with the highest activations appear first in the vector, and genes with the lowest activations appear last. (C) Gene enrichment analysis is then performed on these sorted gene vectors based on gene sets representing KEGG pathways, operons, regulators and GO processes. (D) Finally, the original expression data set is passed through the encoder to associate each of the BN nodes, and their corresponding gene sets, with each growth condition. (E) The first set of seven architectures we have explored with various types of compression and network depth. All architectures have a BN consisting of 50 nodes and an input/output (red bars) matching the number of genes represented in the expression data set being analyzed. (F) The select few additional architectures with their BN layers gradually expanded.

By examining these resulting predicted expression vectors after sorting on expression level (**Fig. 1B**), we associate each BN node with enriched gene sets corresponding to known KEGG pathways, operons, regulators or GO processes (**Fig. 1C**). By then passing the original compendium through the encoder, we can additionally associate each BN node, and by extension the associated gene sets, with the experimental conditions that activate it (**Fig. 1D**).

### Increased network depth amplifies biological inference

We explored three basic network architectures using our established training and inference procedure, including two and three layer encoders/decoders for each (**Fig 1E**; see also Methods section “Construction of the denoising autoencoders with various architectures” for other details on the network construction). Throughout this manuscript, we have adopted the notation:

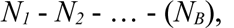

where *N_x_* and *N_B_* are the number of nodes in hidden layer x and the bottleneck layer, respectively, to reference the various architectures with ease. We only specify the number of nodes in each layer for the encoder since the decoder always exactly mirrors this arrangement. The first architecture we chose to have hidden layers with gradual compression based on early work using autoencoders for dimensionality reduction (Hinton and Salakhutdinov, 2006), a choice that has been widely adopted in the deep autoencoder literature (Srivastava *et al*., 2014; Eraslan *et al*., 2019; Lotfollahi *et al*., 2021). We also included network architectures where each layer is of the same width as in the input layer. Finally, we included an architecture which expands the network width by roughly a factor of 2 before compressing the input. We performed our full training and inference process for ensembles of 100 networks for each of the three architectures and two network depths, along with two ensembles for a shallow network architecture to illustrate stability of the process.

There is a clear contrast in gene set recovery between the shallow and deep networks (**Fig. 2A**). In general, deeper networks tend to recover more statistically significant associations with known gene sets (**Fig. 2B**). The differences between architectures of the same depth are often more subtle. This is clearly seen in **Fig. 2B** where the three networks of the same depth, 2000-(50), 4355-(50), and 8000-(50), recover more similar numbers of KEGG pathways, GO processes, and regulators when compared to the sudden jump between the deep and shallow networks.

**Figure 2.**
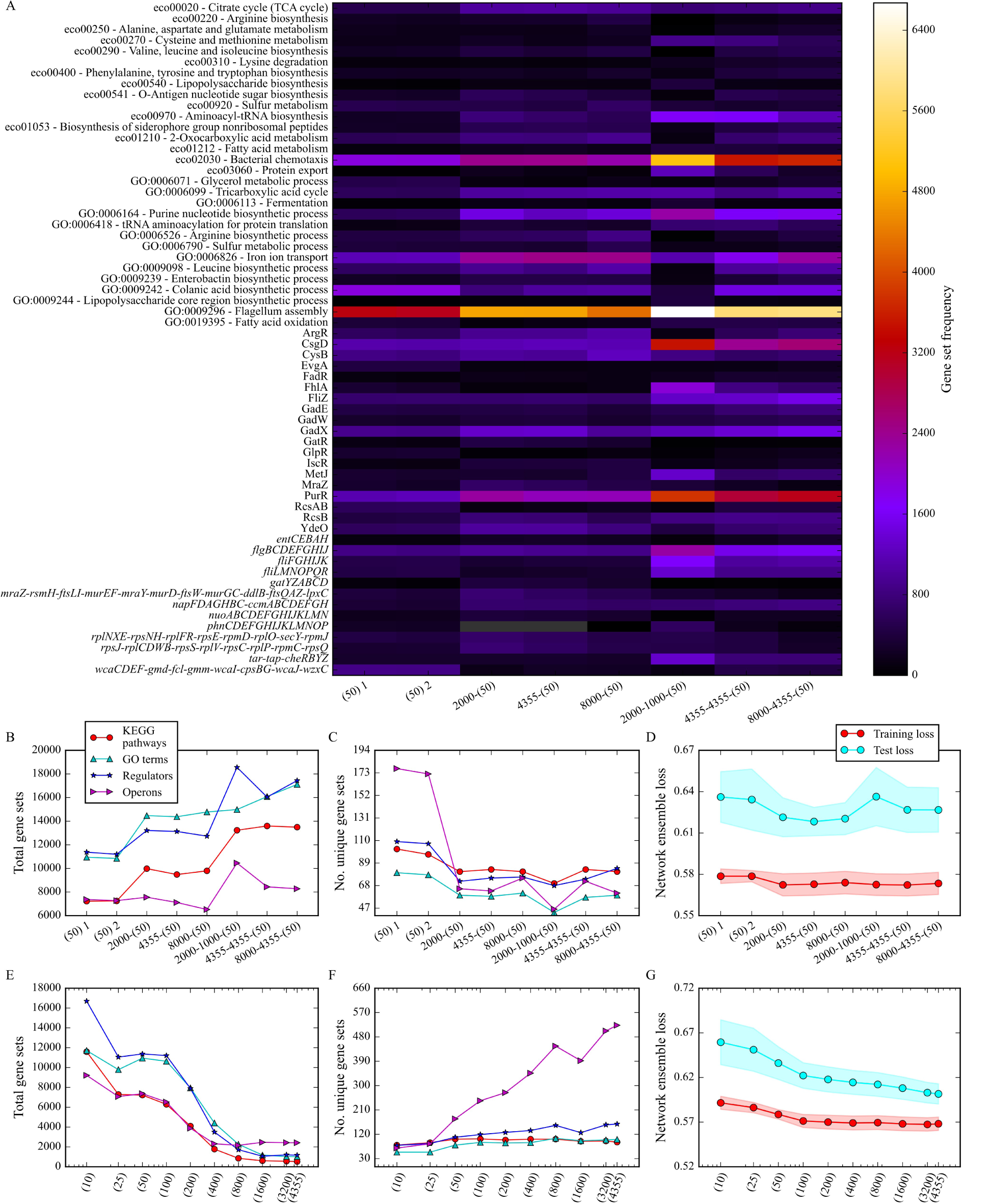
Recovered gene sets for each network architecture. (A) The frequency of all recovered features for each network architecture. The color of each cell represents the frequency of each gene set in the network ensemble for each architecture, while grey denotes no occurrence. Only features which were recovered from at least 3% of bottleneck nodes from the full ensemble of 100 networks for at least a single network architecture are shown. The first two columns are repeats of the shallow DAE ensemble. (B) The total number of recovered gene sets considering even redundant occurrences for each of the seven architectures shown in Fig. 1E. (C) The number of uniquely recovered gene sets in their four respective categories. (D) The average final training and test losses for ensembles of 10 networks for each of the network architectures with single standard deviations shown as envelopes. (E) The total number of recovered gene sets considering even redundant occurrences, in each of the four categories, for the full ensemble of shallow DAEs for various bottleneck sizes. (F) The number of uniquely recovered gene sets in their same four respective categories as E. (G) The average final training and test losses for ensembles of 10 networks for each shallow network compression with single standard deviations shown as envelopes.

A second jump is seen between the networks with single hidden layers between the input and BN, and networks with two layers. Here KEGG pathways show the least overall dependence on other architecture choices apart from network depth. The most dramatic contrast to this is seen in the 2000-1000-(50) networks. These networks appear to learn many more regulators and operons, and slightly less GO processes, when compared to other networks of the same depth. This suggests that the type of gene sets which are recovered is not in general independent of the network architecture, but there is a trend that deep networks recover overall more gene sets.

In contrast, shallow networks recover the most unique gene sets, particularly for operons (**Fig. 2C**). However, these unique gene sets are recovered with a low probability within the ensemble. Similar total numbers of unique gene sets can be recovered from permuted data (**Fig. S8**), suggesting that the number of unique gene sets recovered is not a good proxy for biological inference within the network ensemble. While there were some differences in test loss between networks (**Fig. 2D**), with two-layer encoder/decoder networks showing the lowest average test loss, the differences between network architectures were generally small. Overall, our results indicate that deeper networks tend to more robustly code biological gene sets in their bottleneck nodes.

### A balance between biological inference and the test loss defines optimal compression

Next we have examined how the choice of the number of BN nodes affects gene set inference. This choice, known as the network compression, is often difficult to make just based on prior domain knowledge. To examine how this choice impacts gene set inference, we compared the results of varying the BN layer in the shallow network architecture. We have adjusted the size of the trained network ensembles such that the total number of BN nodes is constant across ensembles (see Methods section “Adjustment of bottleneck layer size”) while spanning several decades of bottleneck layer widths. We believe this choice to hold the number of BN nodes constant is appropriate since each feature can only be found to be associated with a node once per statistical test, and therefore the wider networks have a higher capacity for biological inference.

In terms of inference power, defined as the total number of recovered gene sets, wide networks dramatically lose to the narrower networks (**Fig. 2E**). Although there is a slight upward trend in general in the total inferred unique features as the bottleneck layer is expanded, wider networks are mostly more sensitive to operons (**Fig. 2F**). This suggests narrower networks are forced to learn gene expression patterns associated with broader terms, like GO processes and KEGG pathways, while wider networks are free to code individual transcriptional units, such as operons. There is a plateau in the total number of recovered gene sets between 25 and 100 bottleneck nodes, accompanied by a drop in test loss (**Fig. 2G**), suggesting an optimal trade-off between reconstruction accuracy and biological inference in this range. We have also performed a less-comprehensive local scan over compressions near 50 bottleneck nodes for the deeper network architectures and see a similar trend, although the optimum appears to be closer to 25 BN nodes for the intermediate architecture 4355-(50) (see **Supplemental Fig. S2**).

### Deep networks explain the data with fewer pathways

We next investigated how choice of network architecture impacts which gene sets are reported to be activated or suppressed in each experimental condition. To connect the gene sets associated with each BN node to each condition, we pass each condition into the encoder of each network (**Fig. 1D**), and examine the BN node activations. However, we often observe that a complex combination of both activating and suppressing nodes frequently occurs for a single gene set within a single network for an individual condition. To quantify this aggregate node response as a single number per network per gene set, we used our observation that the relationship between BN node activation and decoder response is approximately linear (**Fig. 3A**). The sum of the decoder outputs when two BN nodes are independently activated is a reasonable approximation to the decoder output when both nodes are activated simultaneously. This linear approximation appears to be accurate across nodes enriched for three distinct gene sets for networks of various depths (**Fig. S3**).

**Figure 3.**
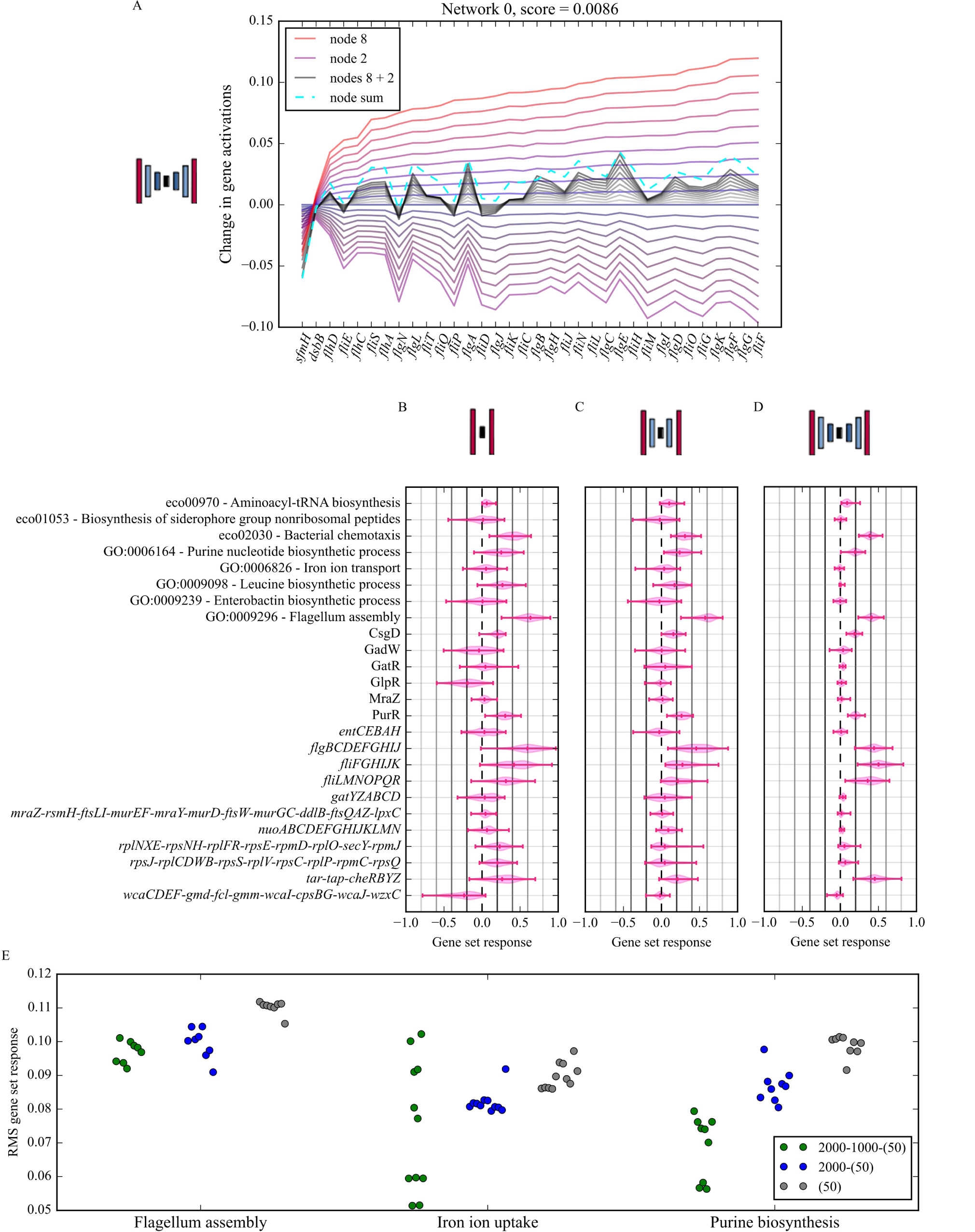
DAE-gene set response to individual experimental conditions. (A) The decoder response for the flagellum assembly GO process gene set when a single activating node from a 2000-1000-(50) network (node 0) is activated gradually from 0 to 1 (red lines of increasing intensity). The purple curves below the x-axis represent the result of the same activations but for a single suppressing BN node (node 5), and the grey curves show the decoder response when both nodes are activated simultaneously. The dashed cyan line is the sum of the maximum decoder responses for both nodes 0 and 5, and the score is the L1-norm between the final grey curve and the cyan line. (B)-(D) The distributions of gene set responses for a subset of all recovered gene sets which show the largest response magnitude when a single randomly-selected experiment which activates flagellum assembly associated nodes is passed through the encoders of the full ensembles for the three architectures. (E) RMS values across all gene sets and networks of the gene set response for the three architectures shown in B-D, computed for sets of experimental conditions which all three network architectures agree upregulate the indicated pathway.

Using this linear approximation, we have defined the gene set response, *ϕ*, to each experimental condition for a single network as

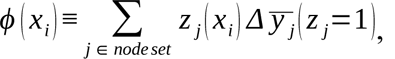

where *z j* (*xi*) is the activation value of BN node j when the experiment, *xi*, is passed through the encoder, and 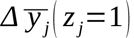 is the average log-fold change in predicted gene expression when the activation of node j is set to 1 with all other BN nodes set to 0, averaged over all genes in the target gene set of interest. This metric represents with a single number the average log-fold change in gene expression for a single gene set as predicted by a single network when a single experimental condition is passed through the encoder.

To investigate how network depth affects gene set response, we randomly chose a single RNA-seq experiment that activates BN nodes associated with the flagellum assembly GO process gene set in all three network architectures (see Methods section “Determination of conditions associated with specific gene sets, and selection of consensus experimental conditions”). We passed this through all 100 networks in each ensemble for each of the selected network architectures. Finally, we computed *ϕ* for all networks for every gene set recovered from the compendium.

Comparing gene set responses between network architectures indicates deeper networks are more selective, with nodes from the deepest networks changing the gene expression for only a few gene sets. The gene sets that show the greatest response according to the deepest networks are eco02030 - bacterial chemotaxis, GO0009296 - flagellum assembly, the operons *flgBCDEFGHIJ*, *fliFGHILK*, *fliLMNOPQR*, and *tar-tap-cheRBYZ* which are all associated with flagellum assembly or motility. Shallow networks also tend to have wider distributions of gene set responses across many pathways, leading to a less clear separation of stimulated gene sets. This selectivity of the deeper networks holds for other gene sets as well. This can be seen in **Fig. 3E** where we have performed a similar process to the above flagellum assembly-associated nodes for iron ion uptake- and purine biosynthesis-associated nodes.

To quantify the selectivity, we have taken the RMS across all networks and gene sets for each architecture, and performed this calculation for a set of consensus experimental conditions which all architectures report activate nodes of the corresponding gene sets (see Methods section “Determination of conditions associated with specific gene sets, and selection of consensus experimental conditions”). On average the deeper networks tend to lower the RMS gene set response and therefore increase selectivity, although this is not true for several select conditions associated with iron ion uptake. This is due to the fact that many of the nodes found in the 2000- 1000-(50) architecture which are associated with iron ion uptake are also associated with down- regulation of flagellum assembly (see **Fig. 4B**) and so inevitably this response is more diverse for these nodes.

**Figure 4.**
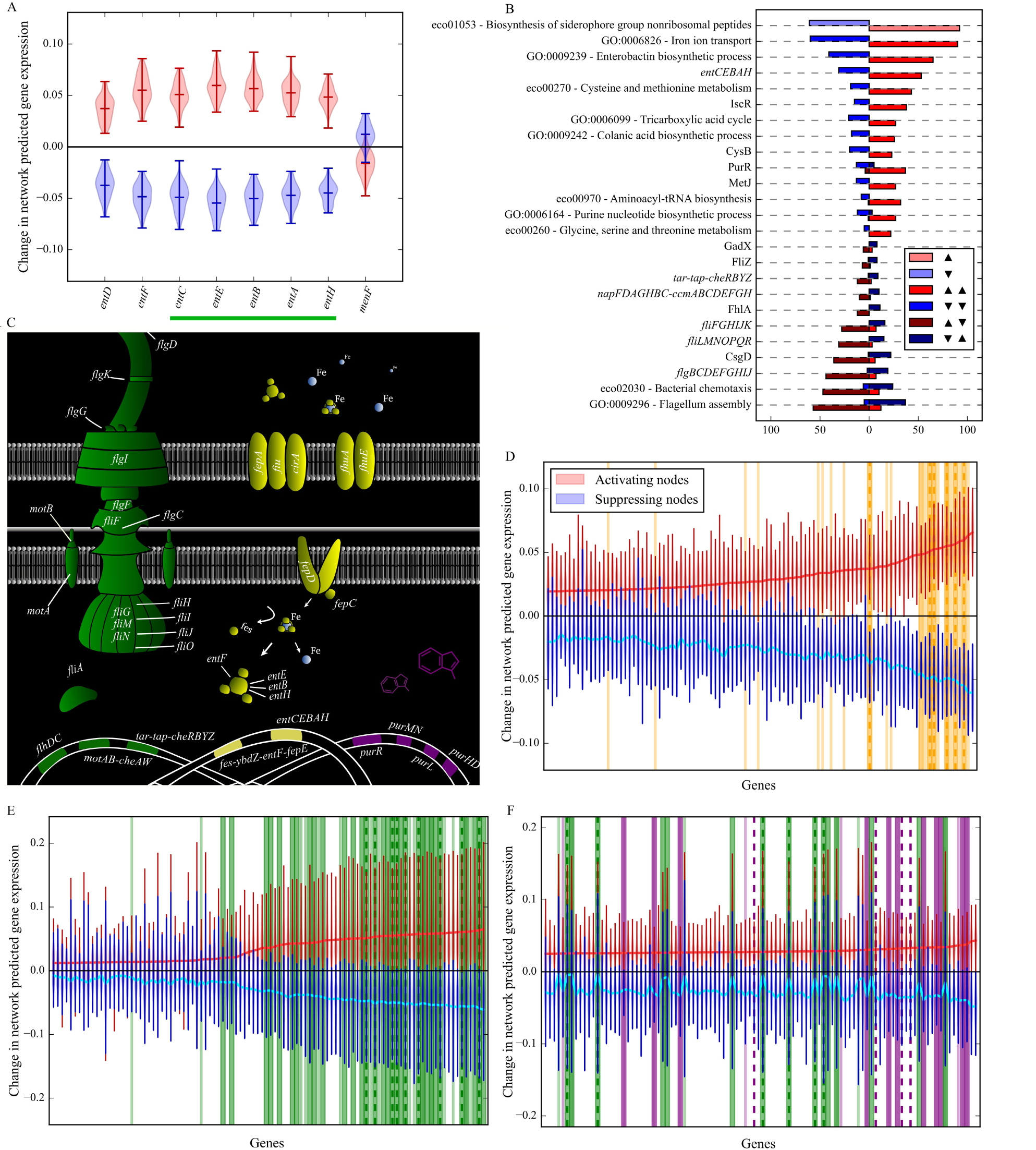
Predicted connection between gene sets by the 2000-1000-(50) network architecture. (A) The distributions of DAE-predicted expression changes for genes in the siderophore biosynthesis KEGG pathway, eco01053. The green bar below indicates that these genes are cooperonic. (B) The number of other gene sets which are also associated with siderophore biosynthesis BN nodes. Whether this siderophore biosynthesis pathway is activated (upward arrow) or suppressed (downward arrow) is indicated by the left set of arrows in the legend. The number of siderophore biosynthesis BN nodes which either activate (light red) or suppress (light blue) this pathway are shown as the first set of bars for comparison. Only gene sets which occur in at least 20% of siderophore biosynthesis BN nodes are shown for brevity. Left-facing bars indicate the number of nodes which suppress (downward arrow) the corresponding gene set on the y-axis, while right-facing bars indicate activation (upward arrow). (C) A simplified version of three biological processes recovered from the PRECISE 2.0 compendium and a rough reconstruction of the locations and functions of the majority of the various proteins involved in the gene sets highlighted in D-F. The colors are chosen to match each category in panels D-F. (D) The top 100 most activated genes, and their corresponding predicted expression changes (red for activating nodes and blue for suppressing nodes), based on the output of the decoder when siderophore biosynthesis nodes are activated. The dark yellow bands indicate genes in the siderophore biosynthesis KEGG pathway, while the lighter, thin bands and vertical, dashed lines indicate genes in the GO processes for iron ion transport and enterobactin biosynthesis, respectively. (E) and (F) show the same quantities, but for nodes associated with the flagellum assembly GO process, and the purine biosynthetic process. The dark green bands, light green, thin bands, and dashed green lines indicate genes in the flagellum assembly GO process, bacterial chemotaxis KEGG pathway, and the *flgBCDEFGHIJ* operon, respectively. Finally, the dark purple bands, light purple, thin bands, and dashed purple lines indicate genes in the purine biosynthesis GO process, the PurR regulon, and the aminoacyl-tRNA biosynthesis KEGG pathway. These gene sets were chosen because they represent the most frequent co-associations (see **Figs. S6B** and **S7B**).

### Autoencoder bottleneck nodes define gene sets beyond their associated enriched terms and predict regulatory connections between disparate processes

An observation from the results of the previous section is that nodes which are associated with particular gene sets are also associated with related gene sets. This is suggested in **Fig. 3D** where the gene sets for bacterial chemotaxis and flagellum assembly show comparable responses even though they overlap by only three genes. This implies that BN nodes represent biological processes beyond the associated enriched terms. Given that the deepest network architecture, 2000-1000-(50), typically produces the most concise response in terms of independent pathway activation (see **Fig. 3D-E**), and also showed the greatest inference power (see **Fig. 2B**), we have chosen this architecture to further examine what biological processes are represented by BN nodes and how different gene sets are grouped together into the same nodes. Additionally, we have focused on the KEGG pathway “eco01053 - Biosynthesis of siderophore group nonribosomal peptides” for ease in examination in part because it contains relatively few genes.

We have examined exactly how each of the siderophore biosynthesis enriched nodes specifically change the expression values for only the genes within this gene set (**Fig. 4A**). Although most of the genes in this set show an increase in expression or are decreased in a similar manner for suppressing nodes, the changes in expression values for b2265, or *menF*, stand out as potentially randomly distributed. The function of *menF*, which is to convert chorismate to isochorismate, is performed by *entC* during enterobactin biosynthesis (Buss *et al*., 2001; Dahm *et al*., 1998) suggesting that these nodes represent this latter process.

To further uncover what process is represented by these siderophore biosynthesis associated nodes, we have examined their other associated gene sets. The additional gene sets with the highest frequencies suggest that the DAEs are learning the mechanism by which iron is actively transported across the outer membrane of *E. coli* via the ferric-siderophore complex (Stintzi *et al*., 2000) (see **Fig. 4B**). A simplified form of this pathway is shown in the middle of **Fig. 4C** where 15 of the top 20 genes predicted by the DAEs (top 100 shown in **Fig. 4D**) are explicitly shown and their roles in the uptake of iron illustrated.

The iron sulfur cluster regulon for the transcription factor IscR also appears in 38 of the 92 siderophore biosynthesis activating nodes, and 15 / 61 of the suppressing nodes. This suggests that these nodes also capture iron-sulfur cluster formation. This is further evidenced by the comparable presence of the cysteine and methionine metabolism GO process gene set as cysteine is often the sulfur source in Fe-S cluster biogenesis (Ayala-Castro *et al*., 2008).

In addition to gene set enrichment, we have also explored which genes show the greatest predicted change in expression when these siderophore biosynthesis nodes are activated to further uncover what these nodes represent biologically. The top 100 genes are shown in **Fig. 4D**. The vast-majority of genes in panel **D** of **Fig. 4** can be trivially connected to iron uptake or iron-sulfur cluster assembly, as even the genes in the *nrdHIEF* operon (explicitly shown on the x-axis as part of this DAE-produced gene set in **Fig. S9**) are regulated by the transcript factors Fur and IscR. Not only does this further support our prior conclusion that these siderophore biosynthesis nodes represent iron uptake as well as Fe-S cluster formation, but this also suggests that BN nodes can be used to generate high-fidelity *de novo* gene sets.

To demonstrate that the gene set reported by the DAEs shown in **Fig. 4D** is indeed upregulated in the top 15 conditions which activate siderophore biosynthesis nodes (see Methods section “Determination of conditions associated with specific gene sets, and selection of consensus experimental conditions”), and downregulate in the bottom 15, we have computed the z-scores from these conditions for the top 50 of the top 100 genes shown in **Fig. 4D** and plotted these values alongside the distributions of z-scores across the entire compendium (**Fig. S9**). Clearly the genes defined by the DAE-generated gene set are up-regulated and down-regulated in the appropriate conditions, with the exceptions of *yddA/B*, further providing evidence for the validity of this gene set generation method.

Apart from identifying common functionality for the genes and gene sets associated with BN nodes, we also note that disparate processes are also often enriched within the same nodes. For instance for the 92 BN nodes associated with *activating* the siderophore biosynthesis pathway, we have found that 57 of these nodes also *suppress* the flagellum assembly GO process (**Fig. 4B**). We also see the converse relationship in a similar proportion: when BN nodes *suppress* the siderophore biosynthesis pathway, they frequently activate flagellum assembly. This implicit relationship between iron uptake and cell motility has been previously described (Inoue *et al*., 2007; Zhang *et al*., 2020), although the underlying mechanism is not known. Purine biosynthesis nodes also often activate flagellum assembly gene sets (**Fig. S7B**), and to a lesser extent the converse is also true (**Fig. S6B**). This pattern is also seen in the top 100 genes for flagellum assembly nodes (**Fig. 4E**) and purine biosynthesis nodes (**Fig. 4F**) where the former is dominated by genes which are clearly linked to the formation and function of flagella, while the latter shows a mix of the two processes. This mixing is also biased towards activating nodes suggesting that flagellum assembly and purine biosynthesis are upregulated together, but not downregulated. Therefore, while BN nodes often represent clearly defined biological processes, they also frequently suggest potential regulatory interactions between processes.

In summary, the gene sets defined by BN nodes are clearly attributed to known mechanisms, suggesting that the gene sets reported by other groups of BN nodes can be interpreted with similar fidelity. Additionally, the enrichment of disparate processes within the same nodes suggests a regulatory connection between these processes, and further suggests that de novo regulatory predictions can be made from BN node groups.

### Pretrained autoencoders reveal gene sets uniquely associated with uropathogenic *E. coli* **infection**

In this final section, we have used the lessons from the previous sections to make gene set predictions on an independent data set including the transcriptome of *E. coli* during a urinary tract infection (UTI) to discover gene sets specifically associated with infection. To accomplish this, we identify BN nodes from the PRECISE 2.0 trained 2000-1000-(50) networks which activate solely during UTI, then use these same nodes for the *de novo* generation of a UTI- specific gene set. We used the transcriptomes of five uropathogenic *E. coli* isolates (HM26, HM27, HM46, HM65, and HM69) grown in lysogeny broth (LB), urine from healthy donors (UR), and directly from participants with urinary tract infections (HUTI) (Subashchandrabose *et al*., 2014). We have passed this dataset into the encoders of the full ensemble of pretrained 100 networks. Out of the 5000 nodes of the whole ensemble, we have identified 14 nodes which only activate in HUTI amongst the three conditions, defining an HUTI-specific group of BN nodes.

To begin investigating the gene set defined by HUTI specific-nodes, we compared it to 4 nodes which activate only in the UR conditions. Interestingly, the gene set defined by the HUTI- specific nodes contains genes which show a negative change in expression for the UR-specific nodes (see **Fig. 5A**, blue distributions), supporting the hypothesis that these genes are specific to the infection process. We next examined the logFC values from the original UTI data set between the UR and HUTI conditions for all 5 strains, and found that the majority of genes are indeed upregulated in HUTI in all isolates (**Fig. 5B**), with the exception of HM65 indicating some degree of strain-specificity.

**Figure 5.**
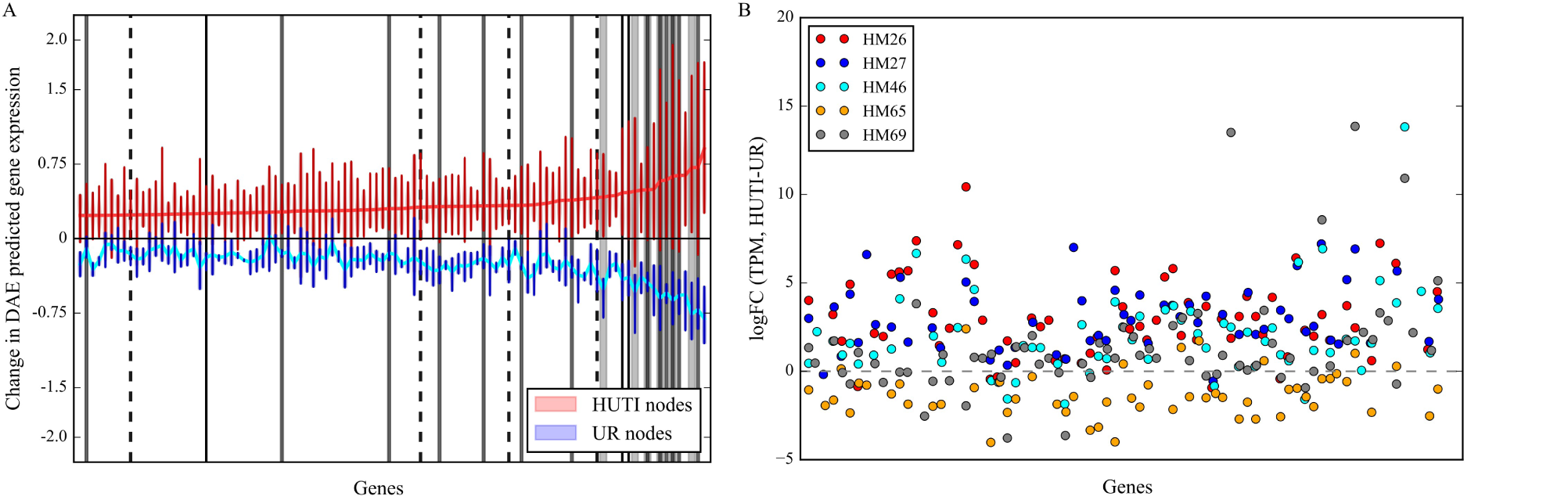
Prediction of the UTI-specific gene set through the ensemble of deep networks with architecture 2000- 1000-(50) pre-trained on the PRECISE 2.0 compendium. (A) The top 100 most activated genes (red line indicating the mean and distributions for the 14 HUTI activating nodes and blue for another 4 nodes found to activate only from UR conditions). The wide, light grey bands indicate genes which correspond to CSPs, while the darker, thin bands indicate genes involved in the Qin prophage. The solid black vertical lines represent genes in the Rac prophage, and the dashed lines genes involved in the DLP12 prophage. (B) The logFCs between HUTI and UR for all five strains from the UTI data set for the same genes shown in A. No marker is shown when the gene is missing in the corresponding strain.

We next examined the functions of these HUTI-specific genes. Amongst this group of upregulated genes, we see six genes encoding cold shock proteins (CSPs), *cspA*, *cspB*, *cspF*, *cspG*, and *cspH*, *cspI*. The *lpxP* gene encoding the cold-inducible lipid A palmitoleoyltransferase was also near the top of the DAE gene list. The HUTI samples examined here were immediately stabilized upon collection using RNAprotect in order to avoid freezing samples in liquid nitrogen (Subashchandrabose *et al*., 2014), making it unlikely that these genes actually represent a response to cold shock. CSPs, particularly CspC and CspE, have been shown to be important for survival in a variety of stress conditions, including infection (Michaux *et al*., 2017; Yair *et al*., 2022). Our results suggest other CSPs may play additional roles in infection.

In addition to the CSPs, many of the genes in our HUTI-specific gene set are encoded on cryptic prophages. Thirteen of the top 100 genes are part of the Qin prophage genes, constituting nearly a third of genes encoded on this island. Other cryptic phage genes include three from the Rac prophage, and four from the DLP12 prophage. Prophages such as Qin are known to be ‘cryptic’ as their machinery for excision and virus production have been rendered inert. The genes encoded on these islands appear to be involved in a range of infection-related processes, including stress tolerance, biofilm formation, and antibiotic resistance (Wang *et al*., 2010)

To investigate the hypothesis that induction of these HUTI-specific genes forms part of a general stress response, we passed the complete PRECISE 2.0 compendium through the encoders of these same 14 HUTI-specific nodes. 12 of the top 30 RNA-seq experiments correspond to strains which have been treated with paraquat at various concentrations to induce oxidative stress. Another 12 experiments correspond to MG1655 strains grown to the antibiotics ciprofloxacin, ceftriaxone, meropenem, or trimethoprim-sulfamethoxazole (Tan *et al*., 2020), again suggesting the HUTI-specific gene set is is stimulated by a variety of stresses.

## Discussion

Here we have explored how network architecture impacts inference from bacterial expression data through DAEs through a pipeline we developed to recover gene sets from ensembles of DAEs. We observed an approximately linear relationship between the activations of the BN nodes and the predicted expression of each gene, and as a consequence we are able to include and interpret deep networks in our analysis, in contrast to previous work (Tan *et al*., 2016, 2017) restricted to shallow networks. Through our analysis, we have found that deeper networks produce more concise gene sets and recover them with a greater frequency. By examining functional categories enriched in these gene sets, we found that BN nodes often recover coordinate regulation of disparate processes. Finally, we demonstrate that deep DAEs can be used to generate *de novo* gene sets, and apply this technique to transcriptomic data from uropathogenic *E. coli* to identify a gene set unique to the infection process consisting of CSPs and prophage genes, potentially involved in response to stressors encountered in the host.

When varying the bottleneck layer width, we found that the test loss of each network ensemble continues to improve even as the training scores converge (**Fig. 2G**). While the wider networks could be finding solutions in which the data is passed from encoder to decoder without alteration, leading to near perfect training and test scores, it appears they are biased towards solutions with more biological significance. Wider networks recovered more operons regardless of network depth (see **Figs. S2B, S2E** and **2F**) and networks with 4355 BN nodes (i.e. no compression) still recover known gene sets in addition to the greatest number of unique operons (**Figs. 2E** and **2F**). This implicit regularization characteristic of deep learning has been described previously for image classification (Barron, 1994; Neyshabur *et al*., 2014).

As a consequence of this ever-improving test loss, we have defined optimal compression as a balance between the improving test loss and the decrease in inference power for wider networks (**Fig. 2E**). However, this decrease appears to be a consequence of our choice to hold the total number of BN nodes constant by varying the number of trained networks in each ensemble. The number of inferred gene sets *per node* decreases as we increase the number of nodes. Alternatively, if we allow the number of networks to vary, we observe that the number of recovered gene sets *per network* only increases as we increase the BN layer (see supplemental **Fig. S1**). Therefore, with wider networks, individual processes are being assigned to individual nodes since these networks have the capacity for this. This distribution of independent processes appears analogous to the concept of disentanglement in variational autoencoders (VAEs), and appears to happen automatically in our DAE pipeline as the BN layer is increased.

Our approach to exploring AE network architectures for gene set inference could be adapted to other types of AEs, such as VAEs which may have advantages in terms of the latent representation of the data learned. A number of studies have explored architecture choices in VAEs, but only by examination of the test score from held-out data on eukaryotic data sets (Lotfollahi *et al*., 2020, 2023; Chow *et al*., 2022; Lotfollahi *et al*., 2021). Our linear decomposition of BN node activations may serve as a complementary alternative to other approaches to AE interpretation, such as training a linear decoder alongside a deep encoder (Lotfollahi *et al*., 2023; Svensson *et al*., 2020).

One advantage of autoencoder analysis is the ability to discover patterns across whole RNA-seq compendia in a hypothesis-free fashion. This contrasts with traditional studies of transcriptional regulators, which are often done in defined inducing conditions. The associations we uncovered indicate several instances of process co-regulation, not easily explained by known regulatory mechanisms. For instance, the regulon of the transcription factor PurR is associated with ∼50% of siderophore biosynthesis nodes, such that these nodes activate both processes (**Fig. 4B**). Another example is seen in the DAE predicted inverse relationship between expression of genes involved in flagellum assembly and those involved in siderophore biosynthesis (**Fig. 4B**), which cannot be easily explained by the activity of the Fur repressor. Such an inverse relationship has been observed previously (Inoue *et al*., 2007; Zhang *et al*., 2020), and was proposed to be mediated by expression of the protein YdiV (Zhang *et al*., 2020). However, we find the *ydiV* gene to be actively suppressed by siderophore biosynthesis nodes, and we also observe this downregulation directly in the data (see **Fig. S9**). Thus, our method serves as a tool for generating hypotheses linking expression of biological processes, and provides a link to the conditions in which connections may be further explored.

## Acknowledgements

This project was funded by the Bavarian State Ministry for Science and the Arts through the research network bayresq.net to LB.

## Methods

### Construction of the denoising autoencoders with various architectures

Autoencoders were implemented using the Python package Keras (Chollet and Others, 2015). All models used sigmoid activation functions at every layer including the bottleneck and output layers. The weights of each layer were randomly initialized based on the Glorot distribution, and bias vectors initialized with zeros. The weight matrices which make up the decoder of each network are tied, i.e. they consist of the transpose of the corresponding weight matrices of each encoder.

### Adjustment of bottleneck layer size

When adjusting the bottleneck layer of each network architecture, the total number of networks in each ensemble is changed such that the total number of bottleneck nodes is constant. For instance, when reducing the number of nodes in the bottleneck from 50 to 25, the number of networks in the ensemble was increased to 200, such that there are still 5000 nodes. For comparison, the case where the number of networks is held constant and the number of BN nodes is allowed to vary was examined, by dividing the results shown in **Fig. 2E** and **F** by the number of networks in each ensemble (see supplemental **Fig. S1**)

### Autoencoder network training and cross-validation

All networks in each ensemble were trained using the Adam optimization algorithm. Data corruption was employed during training such that 10% of entries of each input RNA-seq experiment is randomly set to zero during each training step.

Early stopping was additionally employed to improve generalizability. To do so, each data set was randomly partitioned into an 80% training and a 10% validation set. A 10% test set was separately partitioned to determine optimal training parameters. Local search was performed over all training parameters including the learning rate and batch size using the test score as a metric with batch shuffling enabled.

### Collection of various gene sets

Regulon and operon information was retrieved from RegulonDB (Santos-Zavaleta *et al*., 2019) and KEGG pathways using the KEGG Python module from the bioservices package, in October of 2021. GO processes were extracted from the RefSeq annotation of the complete genome of Escherichia coli str. K-12 substr. MG1655 with accession number NC_000913.

### Statistical tests to associate BN nodes with activation or suppression of specific gene sets

A hypergeometric test was employed to determine gene set enrichment for each bottleneck node vector, including a Benjamini-Hochberg correction for multiple hypothesis testing. Additionally, since many such vectors are examined for representative gene sets (5000 for an ensemble of 100 networks each with 50 bottleneck nodes), a Bonferroni correction was applied such that the expected number of false discoveries is < 1 for a full network ensemble.

Gene set enrichment analysis (GSEA) (Subramanian *et al*., 2005) was performed using each full bottleneck vector as a preranked list, and again controlled for false discoveries. For both statistical tests, we limit the gene set size to at most 40 genes to avoid performing statistical tests on broad categories e.g. “ECO01100 - metabolic pathways.”

### Determination of conditions associated with specific gene sets, and selection of consensus experimental conditions

To determine individual conditions which are associated with up-regulation of specific gene sets, the entire compendium was passed into all 100 encoders of each of the network ensembles. The conditions which activate BN nodes associated with the gene set of interest were then examined. To determine a consensus experimental condition, after performing this previous procedure for all three network architectures, (50), 2000-(50), and 2000-1000-(50), a single condition was randomly selected which appears in the overlap of the three resulting sets of top 50 conditions.

### Gene and protein data acquisition for Fig. 4C

The locations and functions of the various proteins in **Fig. 4C** are taken from (Chang and Liu, 2019; Fujii *et al*., 2017; Johnson *et al*., 2021; Liu and Ochman, 2007) for the genes relating to the flagellum assembly GO process, and (Neumann *et al*., 2018; Zhang *et al*., 2020; Stintzi *et al*., 2000; Kanehisa *et al*., 2023) for genes involved in iron uptake. We have also taken additional supporting information from (Santos-Zavaleta *et al*., 2019).

### Validation on permuted data

To generate the permuted version of the PRECISE 2.0 compendium, gene expression values for each gene were independently shuffled such that the distributions of expression values per gene are preserved, but the gene-gene correlations and condition specificity are destroyed.

## Code availability

Code to run our DAE pipeline as well as the expression data matrix and gene sets used in this study can be directly accessed at https://github.com/BarquistLab/DAE_architecture_exploration.

