## Supplemental material for "Network depth affects inference of gene sets from bacterial transcriptomes using denoising autoencoders"

### Supplemental text to Impact of network architecture on the inference of expression regulation in bacteria using denoising autoencoders

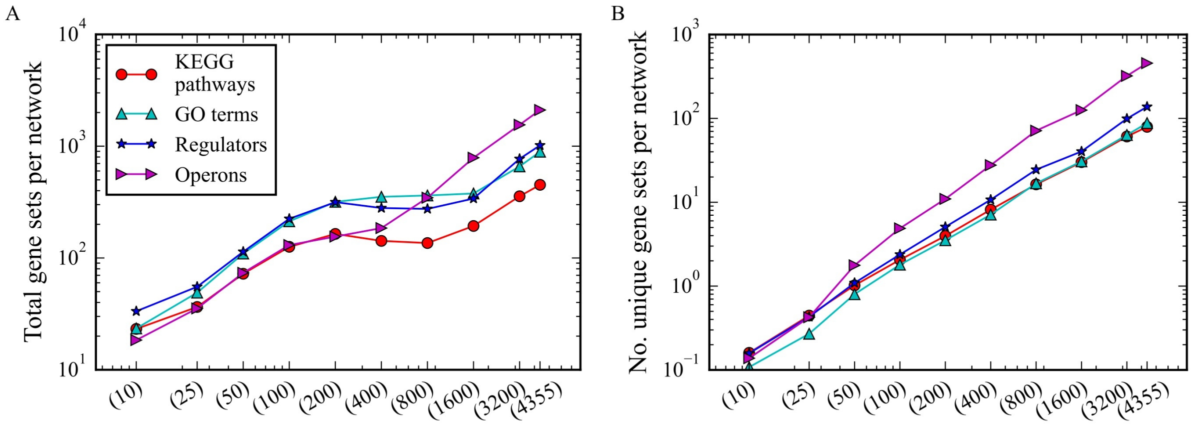


**Figure S1** Recovered gene sets per network for shallow networks with variable BN layers. (A) The total number of recovered gene sets considering even redundant occurrences, divided by the number of networks in each ensemble. (B) The number of uniquely recovered gene sets in their same four respective categories as A divided by the number of networks in each ensemble.

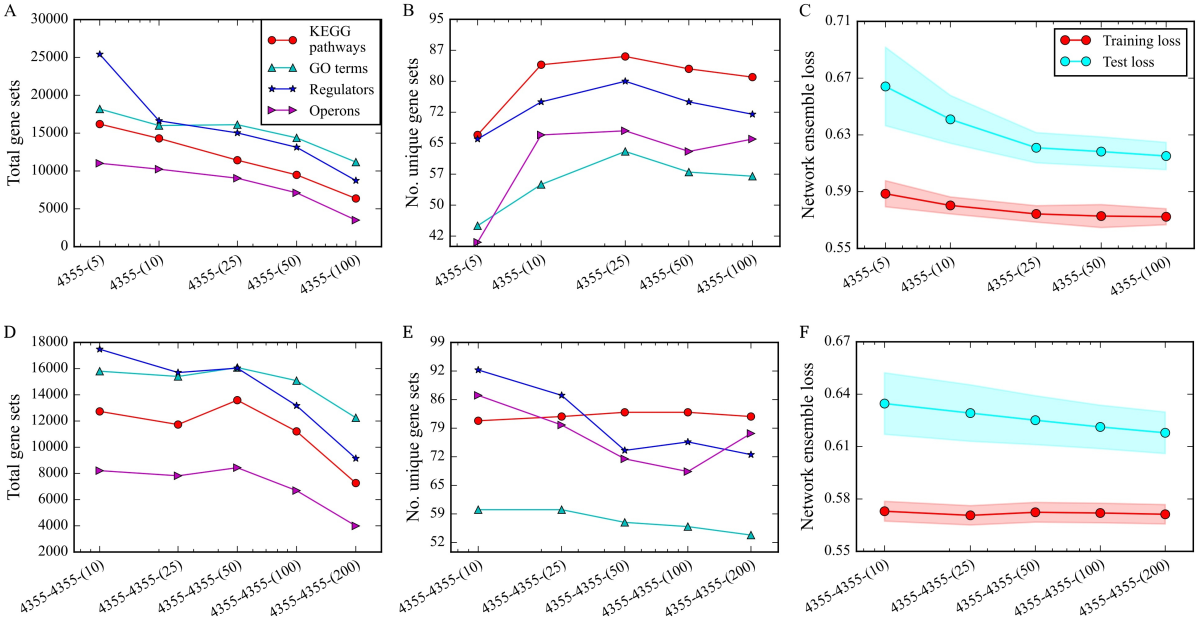

**Figure S2** Recovered gene sets and performances for networks with fixed layer sizes and variable BN layers. (A) Total recovered gene sets considering even redundant occurrences for networks with architecture 4355-($x$), where $x$ is the size of the BN layer. (B) Number of uniquely recovered gene sets in their same four respective categories as A for networks with architecture 4355-($x$). (C) Average final training and test losses for ensembles of 10 networks for each network compression with single standard deviations shown as envelopes. (D-F) Same quantities as in panels A-C but for the 4355-4355-($x$) network architecture.

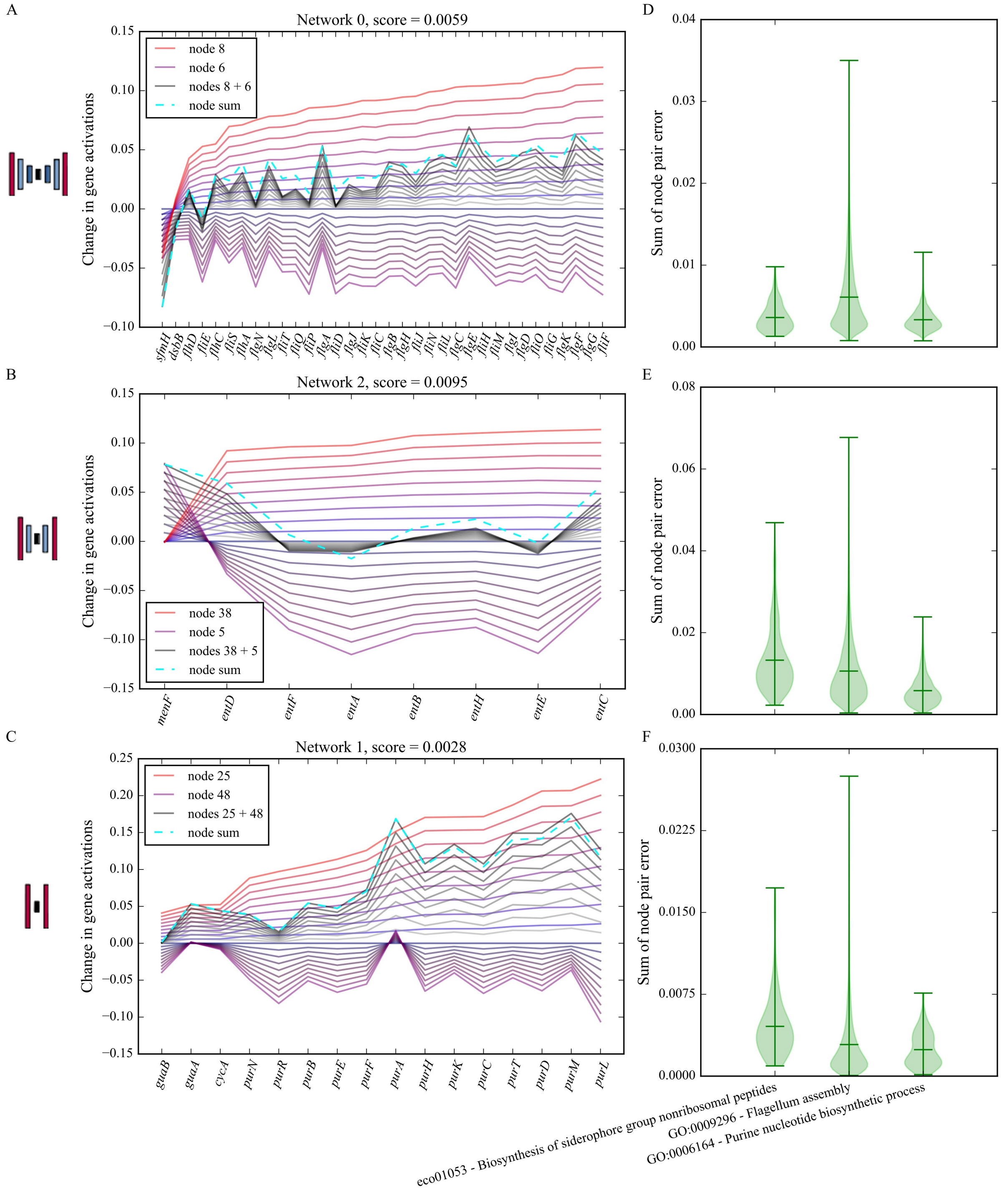


**Figure S3** (A-C) Examples of a typical decoder response for three different gene sets in three different network depths when a single activating node is activated at regular increments from 0 to 1 (red lines of increasing intensity). The purple curves below the x-axis represent the result of the same activations but for a single suppressing BN node, and the grey curves show the decoder response when both nodes are activated simultaneously. The dashed cyan line is the sum of the maximum decoder responses for both nodes. The score at the top of the panel is the L1-norm between the final grey curve and the cyan line. Panel A represents the predicted change in gene expression for the flagellum assembly GO process gene set for a single 2000-1000-(50) network (network 0), B the same for the biosynthesis of siderophore group nonribosomal peptide KEGG pathway gene set from a 2000-(50) network, and C for the purine nucleotide biosynthesis process GO term gene set from a (50) network. (D-F) The distributions of L1-norms, referred to as the sum of node pair errors, for all pairs of activating and suppressing nodes in each ensemble, computed for all three gene sets and all three network depths.

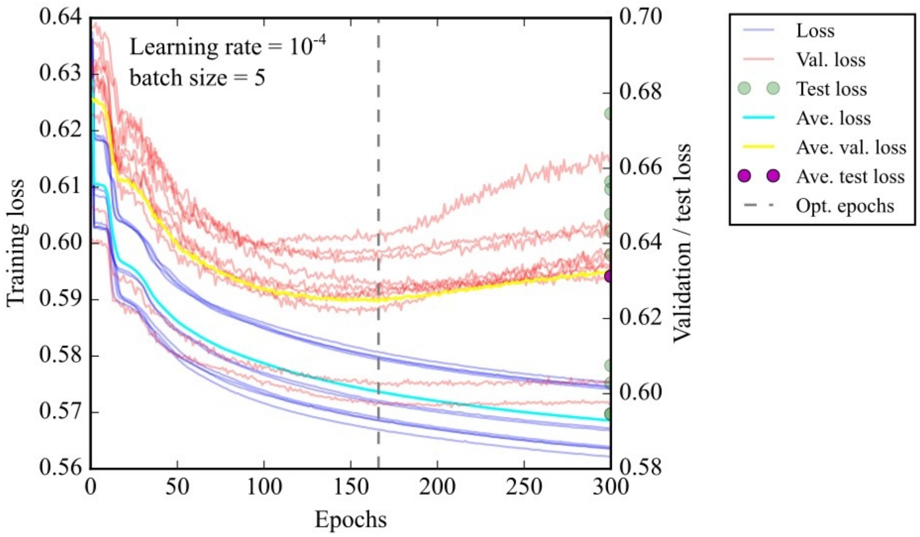


**Figure S4** The training trajectories for an ensemble of 10 networks with architecture 2000-1000-(50). Blue transparent lines indicate the training losses, and red transparent lines the test losses, with each average shown in cyan and yellow, respectively. Additionally, green circular markers indicate the final test loss at the end of each trajectory with the average shown in magenta. The grey dashed line indicates the epoch with the lowest average validation score indicating the point of early stopping.

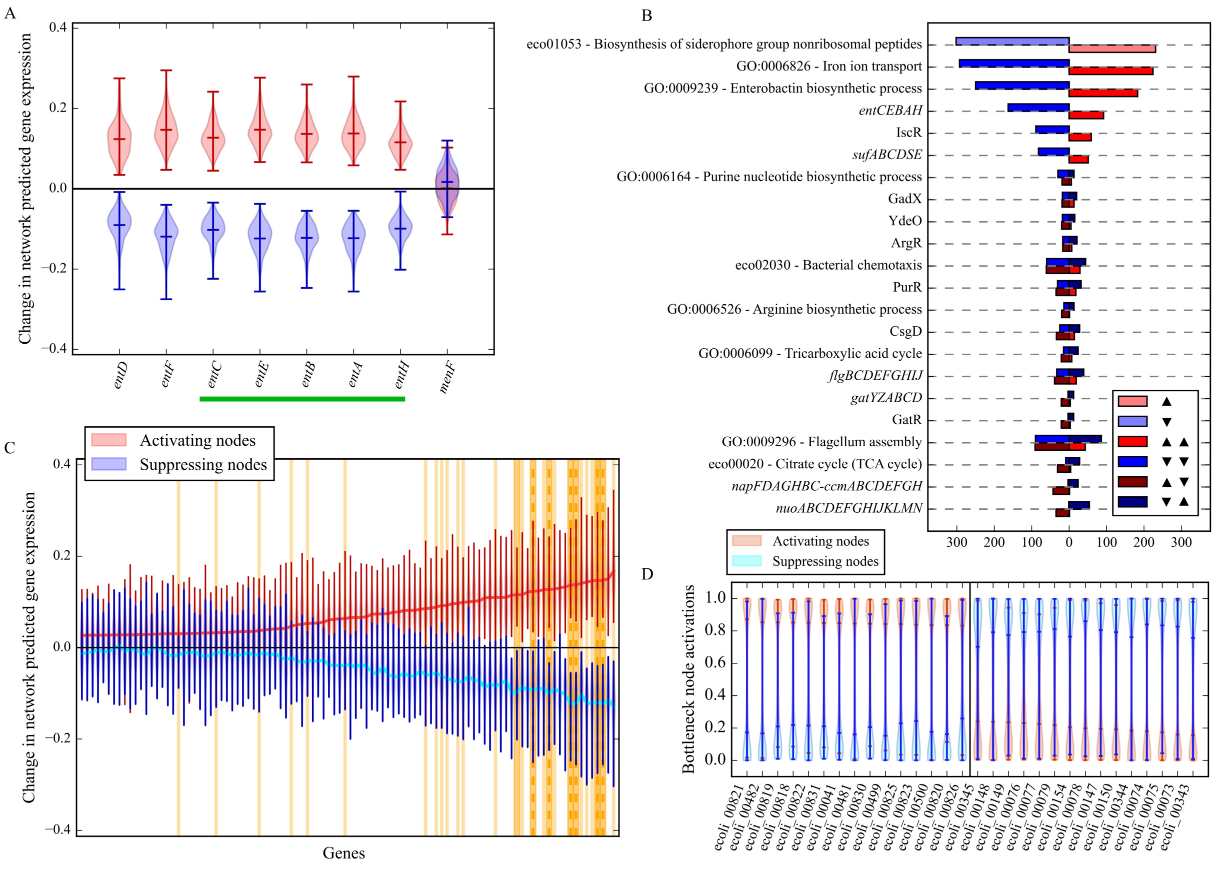


**Figure S5.** Gene activation profiles produced from the first deep network, 2000-(50), for siderophore biosynthesis BN nodes. (A) The distributions of DAE-predicted expression changes for genes in the siderophore biosynthesis KEGG pathway,  eco01053. The green bar below indicates that these genes are cooperonic. (B) The number of other gene sets which are also associated with siderophore biosynthesis BN nodes. Whether this siderophore biosynthesis pathway is activated (upward arrow) or suppressed (downward arrow) is indicated by the left set of arrows in the legend. The number of siderophore biosynthesis BN nodes which either activate (light red) or suppress (light blue) this pathway are shown as the first set of bars for comparison. Only gene sets which occur in at least 20% of siderophore biosynthesis BN nodes are shown for brevity. Left-facing bars indicate the number of nodes which suppress (downward arrow) the corresponding gene set on the y-axis, while right-facing bars indicate activation (upward arrow). (C) The top 100 most activated genes, and their corresponding predicted expression changes (red for activating nodes and blue for suppressing nodes), based on the output of the decoder when siderophore biosynthesis nodes are activated. The dark yellow bands indicate genes in the siderophore biosynthesis KEGG pathway, while the lighter, thin bands and vertical, dashed lines indicate genes in the GO processes for iron ion transport and enterobactin biosynthesis, respectively. (D) The activation distributions for siderophore biosynthesis BN nodes when single experiments of the RNA-seq compendium are passed through the encoder of each network. The distributions are ranked from highest average BN node activation to lowest, with only the top and bottom 15 datasets shown.

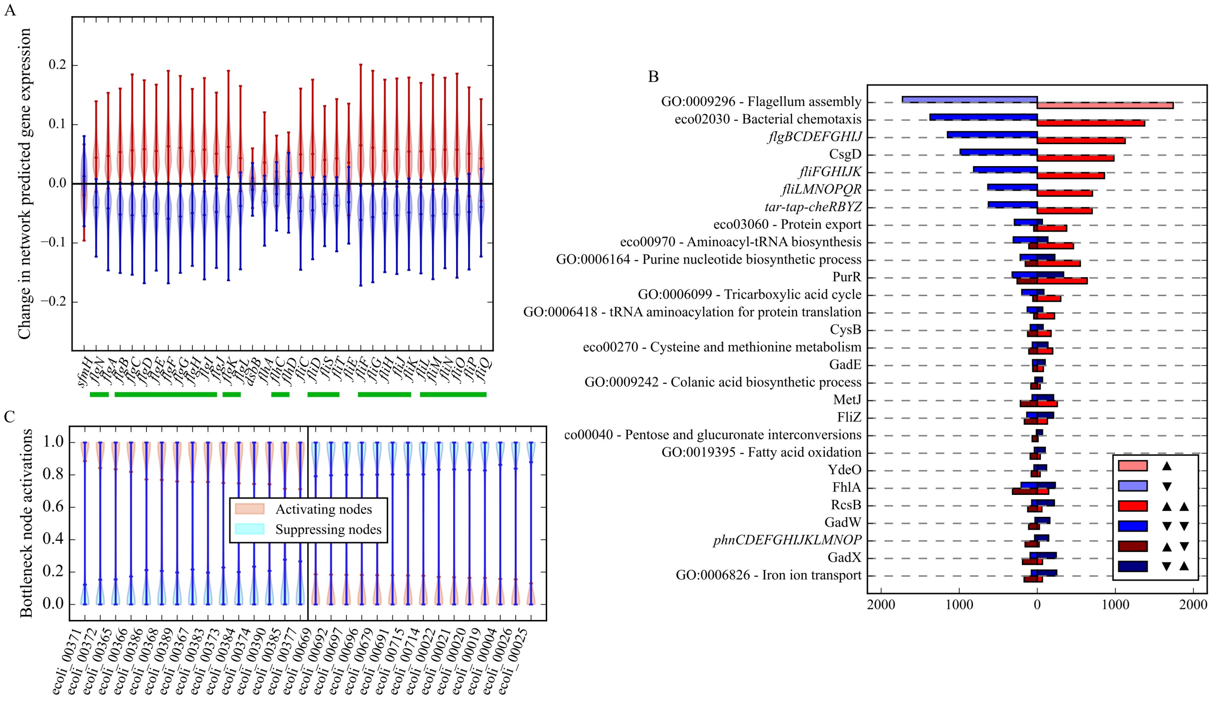

**Figure S6** Gene activation profiles produced from the deep network, 2000-1000-(50), for flagellum assembly BN nodes. (A) The distributions of DAE-predicted expression changes for genes in the "GO:0009296 - Flagellum assembly" gene set. The green bar below indicates that these genes are cooperonic. (B) The number of other gene sets which are also associated with flagellum assembly BN nodes. Whether this flagellum assembly process is activated (upward arrow) or suppressed (downward arrow) is indicated by the left set of arrows in the legend. The number of flagellum assembly BN nodes which either activate (light red) or suppress (light blue) this pathway are shown as the first set of bars for comparison. Only gene sets which occur in at least 20% of flagellum assembly BN nodes are shown for brevity. Left-facing bars indicate the number of nodes which suppress (downward arrow) the corresponding gene set on the y-axis, while right-facing bars indicate activation (upward arrow). (C) The number of other gene sets which were found to be activated or suppressed, for only bottleneck nodes which are also associated with the flagellum assembly GO process (left arrows in the legend). The number of occurrences of the flagellum assembly GO process in both activating (light red) and suppressing (light blue) nodes is shown as the first set of bars for comparison. Only gene sets which occur in at least 20% of the flagellum assembly GO process nodes are shown for brevity. Left-facing bars indicate the number of nodes which suppress (downward arrow) the corresponding gene set, while right-facing bars indicate activation (upward arrow). (D) The activation distributions for flagellum assembly BN nodes when single experiments of the RNA-seq compendium are passed through the encoder of each network. The distributions are ranked from highest average BN node activation to lowest, with only the top and bottom 15 datasets shown.

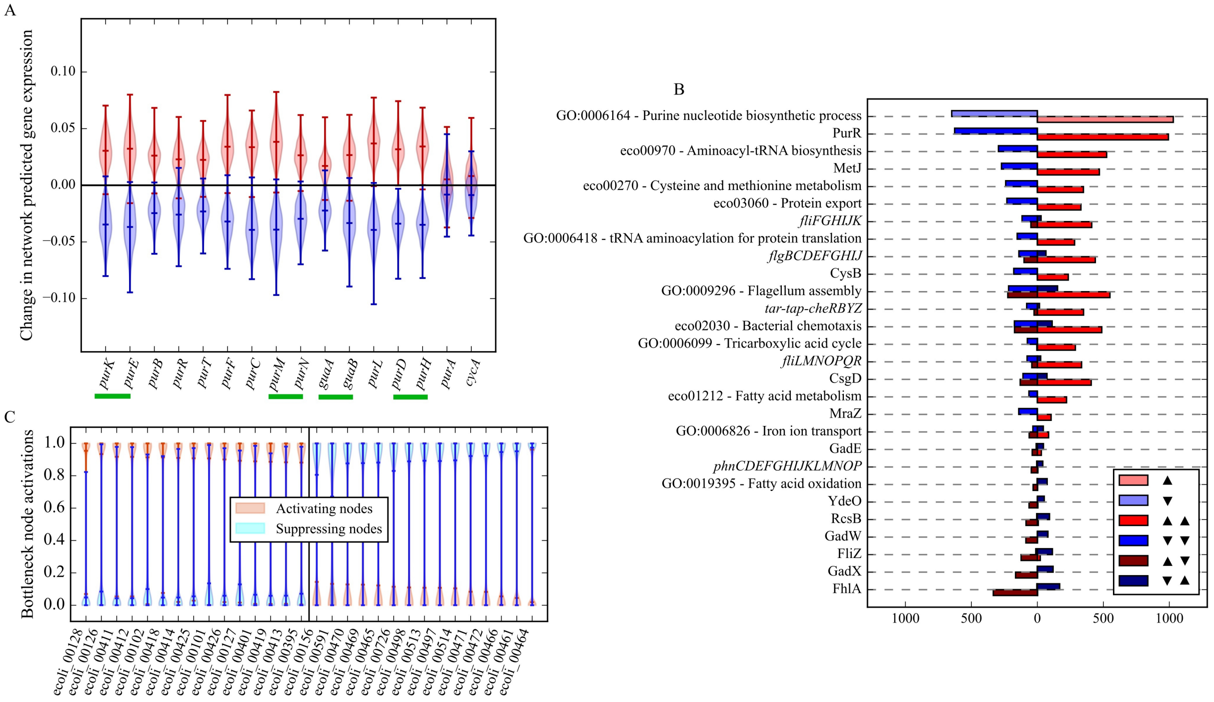


**Figure S7** Gene activation profiles produced from the deep network, 2000-1000-(50), for purine biosynthesis BN nodes. (A) The distributions of DAE-predicted expression changes for genes in the "GO:0006164 - Purine nucleotide biosynthesis process" gene set. The green bar below indicates that these genes are cooperonic. (B) The number of other gene sets which are also associated with purine biosynthesis BN nodes. Whether this purine biosynthesis process is activated (upward arrow) or suppressed (downward arrow) is indicated by the left set of arrows in the legend. The number of purine biosynthesis BN nodes which either activate (light red) or suppress (light blue) this pathway are shown as the first set of bars for comparison. Only gene sets which occur in at least 20% of purine biosynthesis BN nodes are shown for brevity. Left-facing bars indicate the number of nodes which suppress (downward arrow) the corresponding gene set on the y-axis, while right-facing bars indicate activation (upward arrow). (C) The number of other gene sets which were found to be activated or suppressed, for only bottleneck nodes which are also associated with the purine biosynthesis GO process (left arrows in the legend). The number of occurrences of the purine biosynthesis GO process in both activating (light red) and suppressing (light blue) nodes is shown as the first set of bars for comparison. Only gene sets which occur in at least 20% of the purine biosynthesis GO process nodes are shown for brevity. Left-facing bars indicate the number of nodes which suppress (downward arrow) the corresponding gene set, while right-facing bars indicate activation (upward arrow). (D) The activation distributions for purine biosynthesis BN nodes when single experiments of the RNA-seq compendium are passed through the encoder of each network. The distributions are ranked from highest average BN node activation to lowest, with only the top and bottom 15 datasets shown.

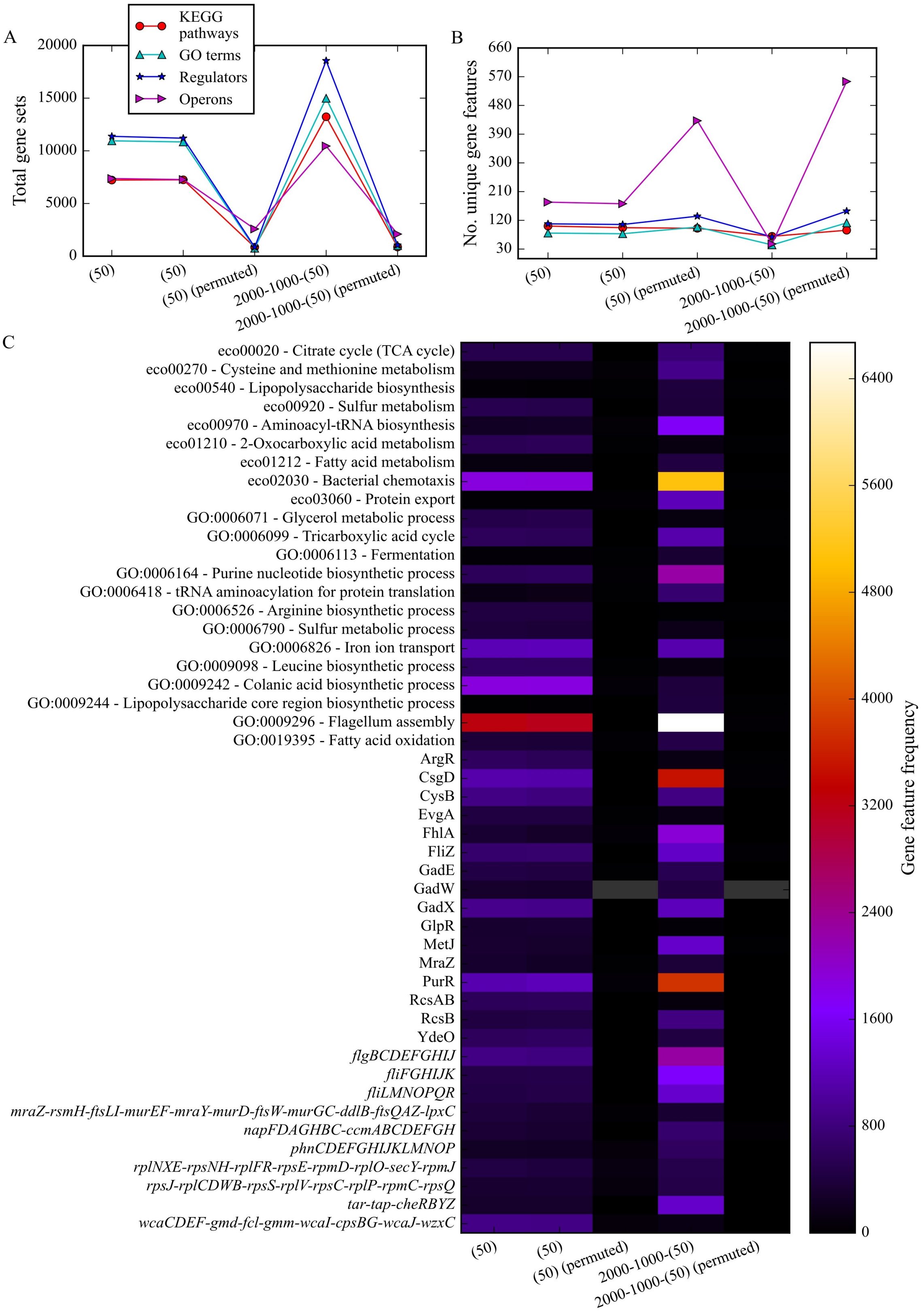
A
KEGG pathways
GO terms
Regulators
Operons
Total gene sets
(50)
(50)
(50) (permuted)
2000-1000-(50)
2000-1000-(50) (permuted)
B
No. unique gene features
C
eco00020 - Citrate cycle (TCA cycle)
eco00270 - Cysteine and methionine metabolism
eco00540 - Lipopolysaccharide biosynthesis
eco00920 - Sulfur metabolism
eco00970 - Aminoacyl-tRNA biosynthesis
eco01210 - 2-Oxocarboxylic acid metabolism
eco01212 - Fatty acid metabolism
eco02030 - Bacterial chemotaxis
eco03060 - Protein export
GO:0006071 - Glycerol metabolic process
GO:0006099 - Tricarboxylic acid cycle
GO:0006113 - Fermentation
GO:0006164 - Purine nucleotide biosynthetic process
GO:0006418 - tRNA aminoacylation for protein translation
GO:0006526 - Arginine biosynthetic process
GO:0006790 - Sulfur metabolic process
GO:0006826 - Iron ion transport
GO:0009098 - Leucine biosynthetic process
GO:0009242 - Colanic acid biosynthetic process
GO:0009244 - Lipopolysaccharide core region biosynthetic process
GO:0009296 - Flagellum assembly
GO:0019395 - Fatty acid oxidation
ArgR
CsgD
CysB
EvgA
FhlA
FliZ
GadE
GadW
GadX
GlpR
MetJ
MraZ
PurR
RcsAB
RcsB
YdeO
flgBCDEFGHIJ
fliFGHIJK
fliLMNOPQR
mraZ-rsmH-ftsLI-murEF-mraY-murD-ftsW-murGC-ddlB-ftsQAZ-lpxC
napFDAGHBC-ccmABCDEFGH
phnCDEFGHIJKLMNOP
rplNXE-rpsNH-rplFR-rpsE-rpmD-rplO-secY-rpmJ
rpsJ-rplCDWB-rpsS-rplV-rpsC-rplP-rpmC-rpsQ
tar-tap-cheRBYZ
wcaCDEF-gmd-fcl-gmm-wcaI-cpsBG-wcaJ-wzxC
Gene feature frequency

**Figure S8** Recovered gene sets for each of the shallow network architectures, and the deep network 2000-1000-(50), from both permuted and unpermuted data. (A) The total number of recovered gene sets considering even redundant occurrences for each of the architectures. (B) The number of uniquely recovered gene sets in their four respective categories. (C) The frequency of all recovered gene sets for each network architecture. The color of each cell represents the frequency of each gene set in the network ensemble for each architecture, while grey denotes no occurrence. Only gene sets which were recovered from at least 3% of bottleneck nodes from the full ensemble of 100 networks for at least a single network architecture are shown. The first two columns are repeats of the shallow DAE ensemble.

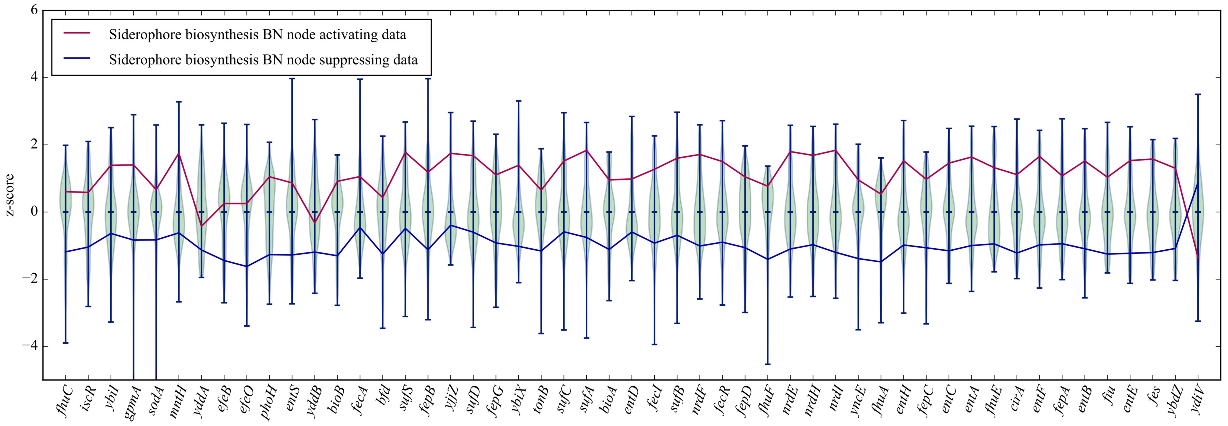


**Figure S9** Siderophore biosynthesis gene set predicted by the 2000-1000-(50) network architecture. The z-scores for the top 50 genes predicted to be most upregulated / downregulated by the DAEs for the top and bottom 15 experiments which activate / suppress the siderophore biosynthesis associated nodes, plotted against the full distributions of z-scores for the entire compendium. The z-scores for the *ydiV* gene are included at the top of this vector, although this is not part of the DAE predicted gene set.
